# Data augmentation for imbalanced blood cell image classification

**DOI:** 10.1101/2022.08.30.505762

**Authors:** Priyanka Rana, Arcot Sowmya, Erik Meijering, Yang Song

## Abstract

Due to progression in cell-cycle or duration of storage, classification of morphological changes in human blood cells is important for correct and effective clinical decisions. Automated classification systems help avoid subjective outcomes and are more efficient. Deep learning and more specifically Convolutional Neural Networks have achieved state-of-the-art performance on various biomedical image classification problems. However, real-world data often suffers from the data imbalance problem, owing to which the trained classifier is biased towards the majority classes and does not perform well on the minority classes. This study presents an imbalanced blood cells classification method that utilises Wasserstein divergence GAN, *mixup* and novel nonlinear *mixup* for data augmentation to achieve oversampling of the minority classes. We also present a minority class focussed sampling strategy, which allows effective representation of minority class samples produced by all three data augmentation techniques and contributes to the classification performance. The method was evaluated on two publicly available datasets of immortalised human T-lymphocyte cells and Red Blood Cells. Classification performance evaluated using F1-score shows that our proposed approach outperforms existing methods on the same datasets.

## Introduction

Blood cells undergo various morphological changes as they progress in their life cycle or undergo the impact of environmental factors. Classification of cell-cycle phases in nucleated blood cells (lymphocytes) is vital for diagnostic and prognostic research studies of pathological conditions and impacts clinical decision making^1^. Likewise, in non-nucleated blood cells (erythrocytes/Red Blood Cells (RBCs)), morphological variations due to storage need to be identified for the prediction of blood quality for life-saving blood transfusions^2^.

Convolutional Neural Networks (CNNs) have exhibited state-of-the-art performances to identify cellular morphologies^2^–4, subcellular localisations^5^ and cell-cycle phases^1,6^. However, the efficacy of CNN based classifiers is logarithmically proportional to the amount of training data^7^. Since data collection in biomedical studies is restricted by the nature of the biological phenomena, data imbalance is a common issue in CNN based classification^7^. In data imbalance settings, some classes have far higher numbers of samples compared to other classes in the dataset. Consequently, samples from majority classes are frequently observed and those from minority classes rarely encountered. The underrepresentation of minority classes causes model training to be heavily biased towards the majority classes, which in turn leads to a significant performance drop of the trained model on minority classes. Furthermore, minority classes often constitute more than half the number of total classes in a biomedical image dataset. Therefore, in order to achieve a model that is applicable in real world settings, it is important for the model to be trained on all the classes equitably.

In existing studies^7,8^, approaches to handle data imbalance can be categorised into parameter-level and data-level. Parameter-level methods alter the learning or decision process by assigning higher weights and often undergo a precarious process of weight determination. Data-level methods include oversampling through various data augmentation techniques to enlarge the minority class image set.

Data augmentation is usually achieved using two types of approaches. The first is the alteration of original images using various image processing techniques such as geometric transformation, colour space augmentation and random erasing, also known as handcrafted methods^9^. Handcrafted methods are efficient, computationally inexpensive and intuitive to apply. However, these techniques take more time to design, as all features need to be mathematically modelled with subjective criteria by humans, which leads to biased decisions and limited improvement. Handcrafted methods could also easily result in overfitting, particularly when some classes contain very few samples. Therefore, techniques such as *mixup*^10^, *Synthetic Minority Oversampling Technique (SMOTE)*^*11*^ *and their variants*^*12*^*–21 have been proposed. mixup* as a data augmentation based model regulariser improves classification performance by linear interpolation of samples to generate synthetic data. Based on *mixup*, Summers et al.^12^ presented multiple nonlinear alternatives, including VH-Mixup (“vertical concat”, “horizontal concat”, and *mixup*)^12^ that combines linear *mixup* with one of their nonlinear methods and performs better than linear *mixup* on benchmark datasets. CutMix^13^ cuts and pastes random patches between the training images and has shown strong classification and localisation ability. MixMatch^14^ is based on a semi-supervised approach that estimates low-entropy labels for unlabelled samples by mixing up labelled and unlabelled images. Balanced-MixUp^15^ performs *mixup* on two images selected using instance-based and class-based sampling simultaneously. Remix^16^ assigns higher weight to the reference image from the minority class and assigns the minority class label to the obtained hybrid image.

The second type of data augmentation uses deep learning based based Generative Adversarial Networks (GANs)^22^ to generate new images by learning the distribution of training data through adversarial learning. Due to their capability of learning data distributions, GANs have been extensively used for image generation, image-to-image translation and image super-resolution^23^. Recently, they have been applied to handle the class imbalance problem as they are able to reproduce the distribution of minority classes^24^–34. On the other hand, training of GANs is complicated with a few common failure modes such as vanishing gradient, mode collapse and unstable training^35^. In order to cope with these problems, Wasserstein GAN (WGAN) was proposed^36^. Advanced versions of WGAN, such as Wasserstein GAN with gradient penalty^37^ (WGAN-GP) and Wasserstein divergence GAN^38^ (WGAN-div), offer more stable optimisation processes and realistic synthetic images. Therefore, recent studies have employed them for various applications such as preprocessing of brain MRI images^39^, image quality improvement of X-ray images^40^ and segmentation of fundus images^41^. However, application of WGAN in biomedical studies to handle data imbalance has not been explored much.

In extreme data imbalance problems, when the number of original samples in the minority classes are a few tens, there is a high chance that synthetic images generated using a single data augmentation approach may appear very similar to each other. Such images often overfit the model, which exhibits poor generalisation ability and does not perform well on unseen data. As demonstrated in our previous study^42^, a combination of WGAN-div and *mixup* generates more diverse samples and helps achieve better classification performance. Extending our previous work^42^, in this study, we propose to include a novel *non*-*linear mixup* along with WGAN-div and *mixup* to further increase the diversity of the training dataset for more robust classification (Fig. 1). We consider that the standard *mixup* generates a synthetic image from two reference images by weighted averaging with limited regularisation effect. In order to increase the regularisation effect, further variation is generated by applying 3D rotation in RGB colour space of the synthetic image obtained from standard *mixup*. In this way, we combine the transformation of pixels in the spatial domain and RGB colour space to achieve better regularisation.

**Figure 1.**
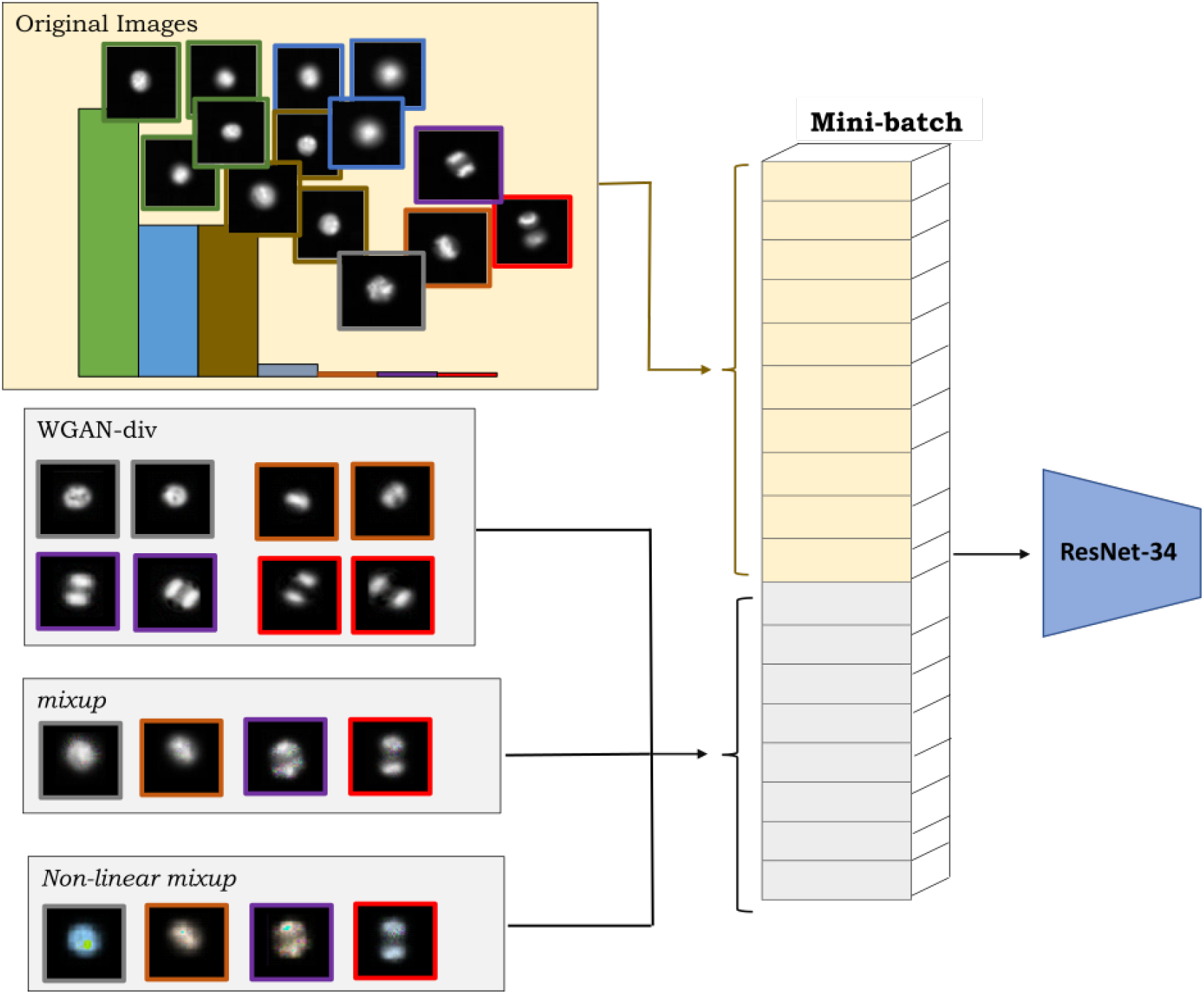
Framework for oversampling of minority classes. In order to enhance the representation of minority classes during training, each mini-batch of original images is supplemented with a batch of only synthetic images of minority classes generated from WGAN-div, *mixup* and the proposed nonlinear *mixup*.

Along with the construction of synthetic data, this study also focusses on designing how these samples are utilised during training. In our previous study^42^, we proposed a minority class focussed sampling approach, which supplements the mini-batch of original images with another mini-batch of synthetic images generated for the minority classes by WGAN-div and *mixup*. In this study, the supplementary mini-batch is formed from the synthetic images generated by WGAN-div, *mixup* and nonlinear *mixup* (Fig. 1). Such minority class focussed sampling allows finetuning of the target distribution to include the minimum number of synthetic samples required, without affecting the representation of original samples.

The proposed method was evaluated on two publicly available datasets: (1) 32, 266 images of immortalised human T-lymphocyte cells (Jurkat cells) (Fig. 2 (a)); and (2) 64, 734 images of RBCs collected from two sites (Fig. 2 (b)) (details in Data description). The results demonstrate that our proposed framework achieves state-of-the-art classification performance for highly imbalanced data distributions on different datasets.

**Figure 2.**
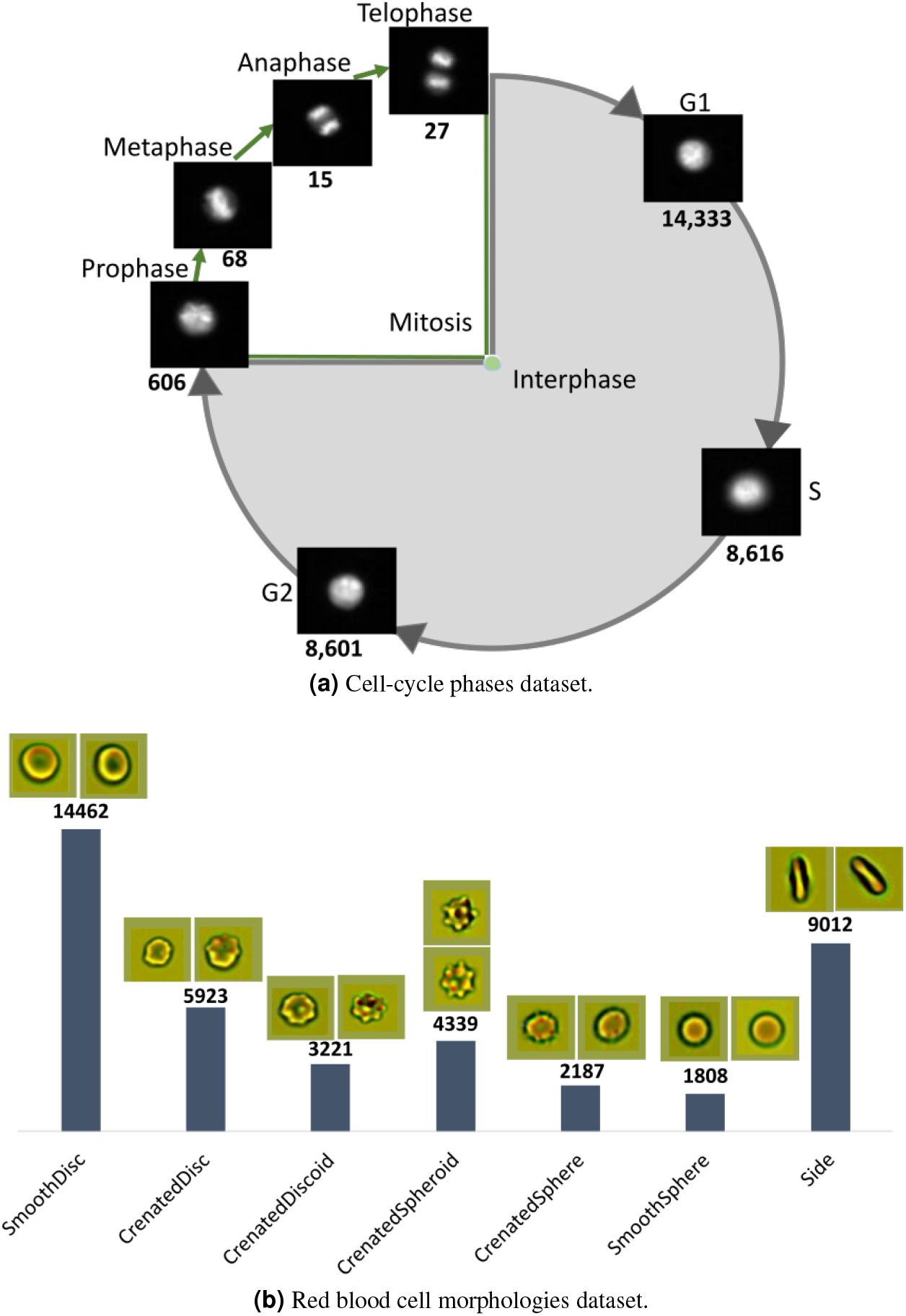
Class distribution of the two datasets used in this study, with the number below each image indicating the number of samples in that class.

## Materials and Methods

### Data description

In this study, experiments were conducted on two public datasets of immortalised human T-lymphocyte cells (Jurkat cells) of different cell-cycle phases and RBCs of different morphologies. Both datasets contain single cell images.

#### Cell-cycle image set

This dataset^1^ consists of 7 classes, representing different phases of the cell-cycle: Interphase (G1, G2, S) and Mitosis (Prophase, Metaphase, Anaphase, Telophase) (Fig. 2 (a)). There are 32,266 pairs of immunofluorescence (IF) and brightfield (BF) microscopic images of Jurkat cells collected by imaging flow cytometry, of which 29, 039 are for training and 3, 227 for testing. Each image of size 66 × 66 pixels is labelled based on two stains: propidium iodine (PI) to quantify each cell’s DNA content and the mitotic protein monoclonal-2 (MPM2) antibody to identify cells in mitotic phases. The number of samples in each class is shown in Fig. 2 (a). The dataset provides separate sets of images for model training and testing, without any overlap of images between training and test sets.

#### Red Blood Cells image set

This dataset^2^ consists of 7 classes: Smooth disc, Crenated disc, Crenated discoid, Crenated spheroid, Side, Crenated sphere and Smooth sphere, and each class represents a morphological state of RBC (Fig. 2 (b)). Images were collected from two sites (Canadian Blood Services and the University Hospital of Geneva)^2^. There are three channels, in which two channels have brightfield images and the third channel has dark-field images. Each image is of size 48 × 48 pixels. Our experiments were performed on 64, 734 labelled brightfield images of RBCs, of which 40, 916 were for training, 14, 764 for validation and 9, 054 for testing. The number of samples in each class is shown in Fig. 2 (b).

### Data augmentation

In our method, we incorporated both handcrafted and deep learning approaches for data augmentation and used three techniques, namely mixup, proposed nonlinear mixup and WGAN-div to generate synthetic samples of minority classes. The primary reason for using more than one data augmentation technique is to increase the diversity of synthetic samples for training.

#### mixup

The mixup method blends two images and generates their hybrid. It regularises the neural network to build a robust model by introducing synthetic images through weighted linear interpolation of input images:

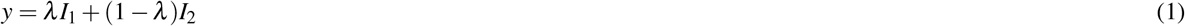

where *I*_1_ and *I*_2_ are two image matrices and *λ* ∈ [0, 1] is the mixup coefficient for each sample pair. Considering each image as a point in 2D space, interpolation along the line joining the two reference images produces a new mixup image on the same line. This method can also be interpreted as the interpolation of each pair of corresponding pixels of two reference images to generate new pixel points, which altogether form the new mixup image.

In this study, mixup samples were generated from a pair of images randomly chosen from the combined set of original and WGAN-div generated images of the same class, therefore the generated sample was labelled as the same class. In order to avoid the resultant *mixup* sample being too similar to the original reference images, *λ* is set to a random value in the range of 0.15 to 0.85 during training.

#### Nonlinear mixup

Standard mixup allows the generation of synthetic points on the line joining the two reference points in the *XY* -plane with limited regularisation effect. Previously, in order to increase the regularisation effect, nonlinear mixup^12^ was proposed to exchange the pixel values of corresponding pixel indices in two reference images in different patterns. So far, nonlinear mixup approaches have been implemented in the spatial domain. Since colour space augmentation^9^ is another successful technique in data augmentation, in this study we utilised the RGB colour space in combination with mixup to increase the regularisation effect.

Consider that the RGB colour space can be geometrically represented as a 3D cube, with red, green and blue components representing each dimension (Fig. 3). Three elements of each pixel of an image (*I*) of size *N*×*N*×3 represents each colour component in the 3D colour space,

**Figure 3.**
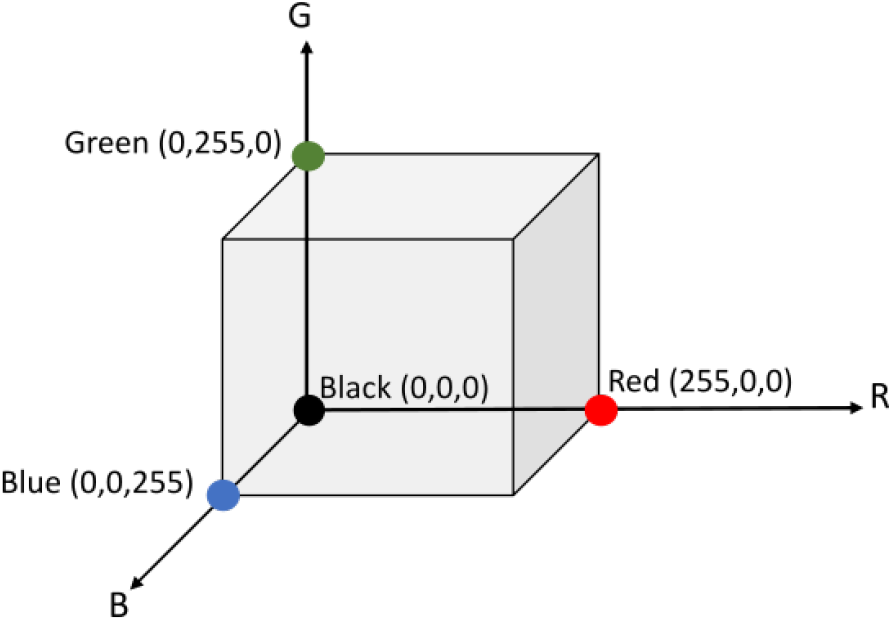
3D cube geometrically representing the RGB colour space.

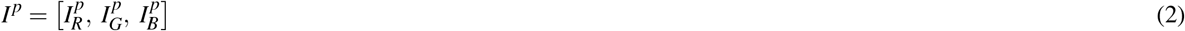

where 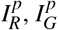 and 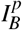 denote the pixel intensities along the red, green and blue colour components (channels) of *I* at pixel *p*.

Since rotation is a widely used transformation, we propose to apply 3D rotation to each pixel of the synthetic image obtained from the standard mixup of two reference images. Furthermore, 3D rotation at an angle *θ* about one of the colour axes allows a controlled tweak in the coordinates and creates new values for the other two colour components (Fig. 4). Subsequently all the rotated pixel values are assembled to generate a new augmented image. In this way, we combine the transformation of pixels in the spatial domain and RGB colour space to achieve better regularisation effect. Using the proposed nonlinear mixup, the synthetic image is consistent with the original spatial distribution of pixels in the mixup image, but with different colour appearance.

**Figure 4.**
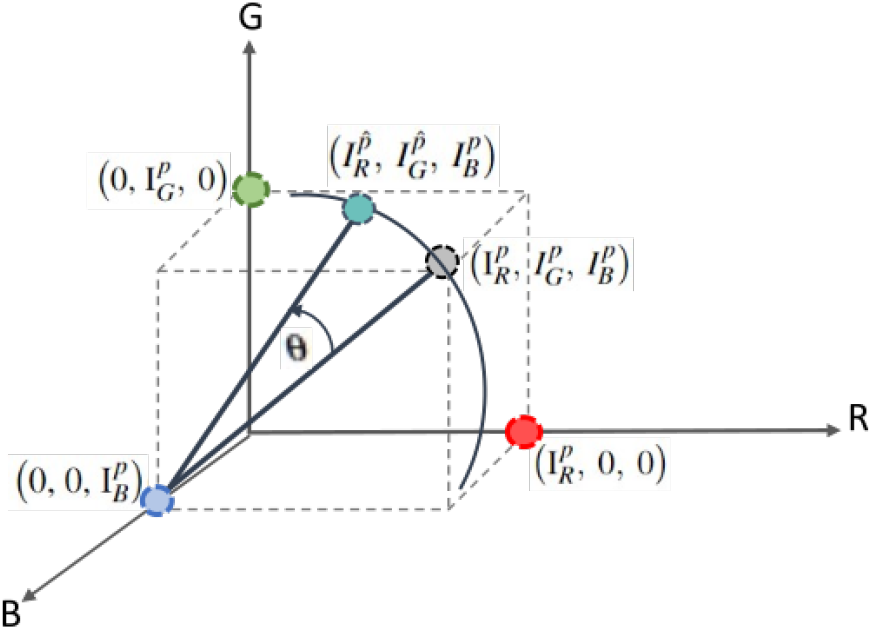
Pixel undergoing 3D rotation.

The proposed nonlinear mixup generates synthetic samples in two steps (Fig. 5):

**Figure 5.**
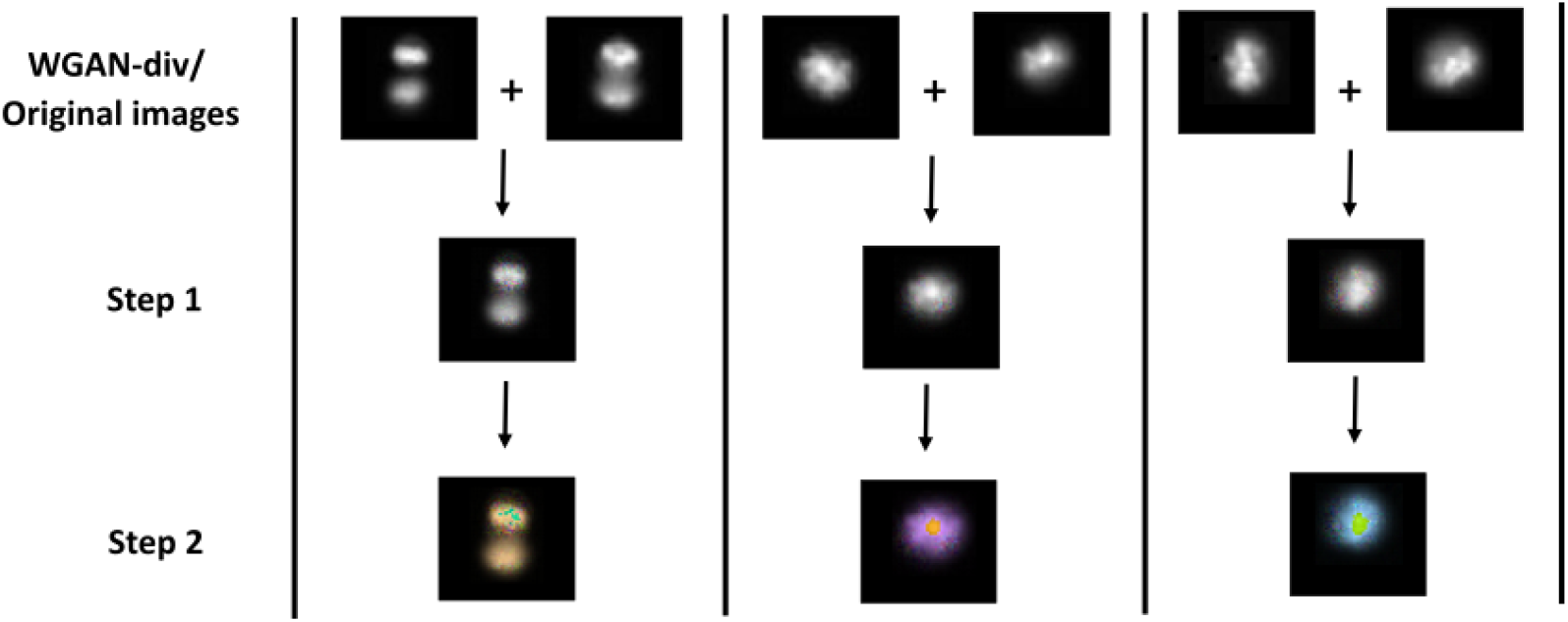
Nonlinear mixup. Step 1: Synthetic image *I*_*mix*_ is generated by applying standard mixup on two reference images from the training set (original and WGAN-div generated images). Step 2: *I*_*mix*_ is further transformed to a nonlinear mixup image by applying 3D rotation to each pixel about one of the axes at angle *θ* in RGB colour space.

1. For two reference images, each with three channels, the standard mixup (Eq. 1) creates a new synthetic image (*I*_*mix*_) in *XY* -plane by applying linear interpolation between corresponding channel images.
2. *I*_*mix*_ from Step 1 is further transformed by applying 3D rotation in RGB colour space. Specifically, each pixel undergoes 3D rotation about one of the colour axes at angle *θ* and gets transformed to a new pixel value. For instance, as shown in Fig. 4, a pixel in 3D colour space represented as 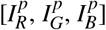 is rotated about the blue axis *B* at angle *θ* using rotation matrix *Rot*_*B*_. A new pixel value is created on a circle with centre 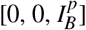 and radius 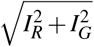 as:

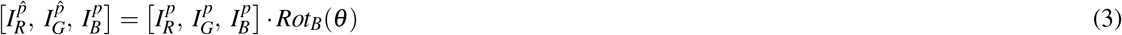

The rotation matrices for 3D rotation about *R, G* and *B* axes are *Rot*_*R*_, *Rot*_*G*_, *Rot*_*B*_ respectively:

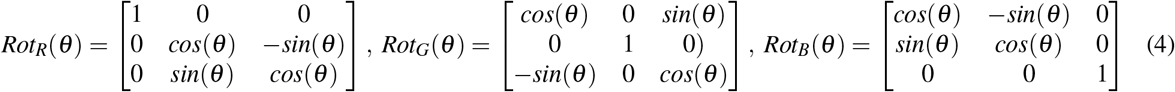

Following Eq. 3 for all pixels, we generate transformed pixel values, which are then assembled to produce the nonlinear mixup image (Fig. 5). We note that for gray scale images, the pixel value in all three channels is the same (*I*_*R*_ = *I*_*G*_ = *I*_*B*_). In this study, *θ* is randomly chosen in the range of 0^◦^ to 360^◦^ and the colour axis to apply 3D rotation is chosen randomly whenever nonlinear mixup is performed.

The combination of transformations in the spatial domain and colour space creates another augmented synthetic image which is still in the spatial domain of the original images, but yields a different colour appearance. Subsequently, the new image generated using nonlinear mixup contributes to the diversity of the training set, improves the regularisation effect and yields improved performance.

#### WGAN-div

Other than handcrafted approaches, data augmentation can also be performed using a deep learning based GAN model, which learns the distribution of a dataset and generates synthetic samples from random noise. A GAN models complex data distributions through joint optimisation of two networks: the Generator (G) and Discriminator (D). The G network generates synthetic/fake data and the D network decides whether the generated samples are real or fake. Both networks are trained alternatively, with the primary goal of maximising the probability of classifying real images as real and generated samples as synthetic/fake. The GAN is well-trained when the generator learns to generate data samples that are as diverse as the original data distribution and can fool the discriminator into accepting them as real.

Training of GAN is coupled with a few common failure modes such as vanishing gradient (discriminator reaches perfect optimisation that does not provide any valuable information for the generator to get better), mode collapse (generator produces limited similar samples) and unstable training^35^. The loss function utilised for GAN training is one of the primary reasons for the training related issues in GAN. As an improved GAN, WGAN^36^ employs a new loss function based on the Wasserstein distance (W-Dis) instead of the Jensen-Shanon Divergence (the standard objective function of GAN) to compute the loss. W-Dis estimates how easy it is to distinguish between synthetic and real images, giving rise to valuable gradients. The objective function of WGAN is:

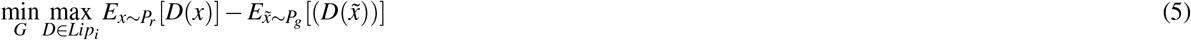

where 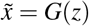 is a generated sample with (*z* ∼ *P*_*z*_) as random vector. *P*_*r*_ represents the distribution of real data, *P*_*g*_ is the distribution of fake data generated by G and *Lip*_*i*_ is the Lipschitz constraint.

WGAN offers more stabilised training than GAN, however the computation of W-Dis requires a 1-Lipschitz constraint, which is a strict constraint. WGAN explicitly clips the weight of the critic *D* within a compact space to maintain the Lipschitz constraint, which often creates convergence issues. Therefore, WGAN-GP was proposed to improve WGAN that uses gradient penalty to enforce the Lipschitz constraint. As an improved WGAN, Wu et al.^38^ proposed Wasserstein divergence GANs (WGAN-div) which uses Wasserstein-divergence (W-div) to compute the objective function instead of W-Dis. WGAN-div approximates W-div through optimisation, without the 1-Lipschitz constraint and offers more stabilised training. The objective function of WGAN-div is:

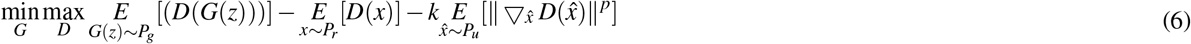

where *p* and *k* are hyperparameters that control the gradient, *P*_*u*_ is a Radon probability measure, *z* is random noise and 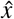 is sampled as a linear combination of real and fake data points. Additionally, WGAN replaces GAN’s discriminator model with a critic that outputs a scalar score which reflects the quality of the generated sample. The lower the critic loss, the higher the quality of the generated image.

In order to handle data imbalance, this study utilised the WGAN-div model to generate synthetic samples for minority classes. A WGAN-div model was trained on the original training set images of each minority class in both datasets. In the cell-cycle phases dataset, the minority classes are Anaphase, Prophase, Metaphase and Telophase, while in the RBCs dataset, the minority classes are Crenated disc, Crenated discoid, Crenated spheroid, Crenated sphere and Smooth sphere.

For the cell-cycle phases dataset, images from each minority class were augmented using affine transformations, perspective transformations, contrast changes and Gaussian noise with diverse parameters to increment the image number to 800 - 1000 for training of the WGAN-div model. Since minority class samples in RBC morphologies dataset are in the thousands, no augmentation was performed. WGAN-div training was guided by the critic loss value. As the absolute critic loss value steadily reaches a state from where it does not decrease further or starts increasing, it was considered a convergence point and the corresponding model checkpoint was used to generate the synthetic samples. Consequently, 3, 000 samples for Telophase, three sets of 6, 500 samples each for Anaphase, Prophase and Metaphase and 4, 000 samples for each minority class in RBC morphologies dataset were used as the augmented images.

### Minority class focussed sampling

During training with an imbalanced dataset, a standard sampling approach would compile a batch with very few images from minority classes, therefore the model training would be heavily biased towards the majority classes. Even with data augmentation, the participation of minority classes during training remains limited. Therefore, in order to ensure balanced representation of samples from all three data augmentation techniques and classes in the dataset, we utilise our previously proposed approach, namely minority class focussed sampling^42^. Under this approach, the mini-batch of original images is supplemented by another mini-batch of only minority class synthetic samples generated by utilised data augmentation techniques. In the current study, supplementary mini-batches were composed of synthetic samples generated by WGAN-div, *mixup* and the proposed nonlinear *mixup*. The number of synthetic samples included can be finetuned using hyperparameter *n* (Fig. 1). Since the synthetic images are not merged with the original samples and are included as a supplementary batch, the representation of original samples during training is not affected.

Estimation of sample distribution ratio (proportion of majority to minority samples) during oversampling is a crucial step for effective and efficient training of the classification model. As a standard practice of performing oversampling, the sample distribution ratio is preset, which decides the number of samples to be added to the original set for achieving the desired ratio^43^. However, in our experiments we observed that the number of synthetic samples added does not necessarily need to satisfy a set ratio. In order to save computational cost, it is important to identify the least effective number of samples for the best possible performance. Therefore, we designed a sampling approach which allows finetuning of the number of synthetic samples added in each batch. In this way, it is not necessary to preset the sample distribution ratio and the number of samples can be finetuned from 1 to *n* for each class during an iteration, where *n* = 1 adds four images (2 WGAN-div + 1 *mixup* + 1 nonlinear *mixup*) of the minority class to the batch. Accordingly, the batch size of original images was adjusted to maintain the overall (original samples + synthetic samples) batch size of 2^*m*^, where *m* is usually *>* 4. The proposed approach reduces computational cost and allows the inclusion of the minimum number of synthetic samples, without affecting the representation of the original samples.

### Experimental setup and implementation

The proposed method to handle data imbalance was evaluated on two datasets: cell-cycle phases and RBC morphologies dataset. The original images from the training set were used to train four WGAN-div models in the cell-cycle phases dataset and five WGAN-div models in the RBC morphologies dataset, one for each minority class. WGAN-div generated images along with the original images were further used to construct *mixup* and nonlinear *mixup* samples, which were then used for training along with the original images using the minority class focussed sampling approach.

We explored alternatives for WGAN-div such as WGAN-GP, WGAN and GAN. All the models utilised default model architectures and optimiser for model updates as proposed in their respective original studies. Accordingly, WGAN-div, WGAN-GP and GAN used the Adam^44^ optimiser, while WGAN used the RMSprop^45^ optimiser. Batch size was set to 16. Hyperparameter *n*_*critic*_ controls the number of times critic is updated for each update to the generator model, and it was set to 4 for all the minority classes of both datasets except Prophase for which *n*_*critic*_ was set to 10. Likewise, the learning rate for all the minority classes of both datasets was set to 0.0002 except Prophase for which the learning rate was 0.00002 (see Discussion section for a justification of parameter choices). Hyperparameters *p* and *k* for WGAN-div were set to their default values of 6 and 2 respectively^38^. The clipping threshold for WGAN-GP was set to 0.01. The learning scheduler and other hyper-parameter initial settings (batch size, decay, learning rate) were the same for all the experiments. We note that the Frechet Inception Distance score^46^ (FID scores) vs epoch was plotted for GAN models (see Supplementary Fig. S3, S4), and as the FID score converges to a constant value, the corresponding checkpoint was used to generate the GAN images.

For the classification of cell-cycle phases dataset, the available training set was divided as follows: 80% for training, and 20% for internal validation, during 5-fold stratified cross-validation. Subsequently, five models were obtained after training on each set of training and validation images. The best performing model was selected and used to evaluate the available test set images. The classification model was trained by finetuning the ImageNet pretrained ResNet-34^47^ architecture with cross-entropy loss. The mini-batch size was 32, where the number of original samples was 8 and *n* was set to 2 for Anaphase and Metaphase, and 1 for Prophase and Telophase. The weight parameters were updated using the momentum Stochastic gradient descent method with a momentum parameter of 0.9 and weight decay of 0.0001 over 200 epochs. The initial learning rate was 0.03, StepLR was used to schedule the learning rate with gamma and the step size was set to 0.1 and 40 respectively. For RBC morphologies dataset, a validation image set has been provided along with the training and test sets, therefore cross-validation was not performed. The classification model was trained by finetuning the ImageNet pretrained ResNet-50^47^ architecture with cross-entropy loss, as ResNet-34 underwent overfitting with RBC images. The mini-batch size per iteration was 32, where the number of original samples was 12 and *n* was set to 1 for all the minority classes. The weight parameters were updated using the momentum Stochastic gradient descent method with a momentum parameter of 0.9 and weight decay of 0.0001 over 70 epochs. The initial learning rate was 0.003, StepLR was used to schedule the learning rate with gamma and the step size was set to 0.1 and 30 respectively.

Images were resized to 64 × 64 pixels for classification using bilinear interpolation for RBC morphologies dataset and cropping for cell-cycle phases dataset. ReLU was utilised as the activation function. Random combinations of image transformations such as horizontal/vertical/left/right flips, rotation at 90^◦^, 180^◦^, 270^◦^ and brightness of images in the range of 50-150% of the original pixel value were also applied. We used macro F1-score for performance evaluation of the proposed method (Table 1 and 2), which is the arithmetic mean of the F1-scores of all the classes and captures per-class accuracy. Macro F1-score is a more suitable metric than accuracy in imbalanced data settings, as the latter indicates the classification performance of the majority classes while macro F1-score reflects the performance of all the classes.

**Table 1.** Comparison with previous studies.

| CELL-CYCLE PHASES DATASET |  |  |
| --- | --- | --- |
| Study | Accuracy | Macro F1 |
| Eulenberg et al.<sup>1</sup> | 0.790 | 0.632 |
| Jin et al.<sup>6</sup> | 0.820 | 0.760 |
| Base model | 0.821 | 0.545 |
| Rana et al.<sup>42</sup> | 0.850 | 0.860 |
| Our method | **0.856** | **0.884** |

**Table 1.** Comparison with previous studies.
| RBC DATASET |  |  |
| --- | --- | --- |
| Study | Accuracy | Macro F1 |
| Doan et al.<sup>2</sup> | 0.765 | 0.767 |
| Base model | 0.759 | 0.760 |
| Our method | **0.820** | **0.800** |

**Table 2.** Ablation study of each component (WGAN-div, *mixup*, nonlinear *mixup* and minority class focussed sampling). Performance comparison between different GAN models, sampling approaches and nonlinear *mixup*. p-values smaller than 0.01 implies the higher statistical significance of the proposed method than the compared methods.

| Data Augmentation Method | CELL-CYCLE PHASES DATASET |  |  | RBC MORPHOLOGIES DATASET |  |  |
| --- | --- | --- | --- | --- | --- | --- |
|  | Accuracy | Macro F1 | p-value | Accuracy | Macro F1 | p-value |
| WGAN-div + *mixup* + nonlinear *mixup* | **0.856** | **0.884** |  | **0.820** | **0.800** |  |
| WGAN-GP + *mixup* + nonlinear *mixup* | 0.852 | 0.860 | 1.5e-04 | 0.814 | 0.794 | 2.8e-7 |
| WGAN + *mixup* + nonlinear *mixup* | 0.842 | 0.810 | 7.2e-04 | 0.778 | 0.789 | 1.4e-9 |
| GAN + *mixup* + nonlinear *mixup* | 0.840 | 0.802 | 1.2e-05 | 0.777 | 0.775 | 3.2e-9 |
| WGAN-div + *mixup* + nonlinear *mixup*<sup>12</sup> | 0.845 | 0.800 | 1.8e-08 | 0.780 | 0.771 | 1.6e-10 |
| WGAN-div + *mixup* | 0.854 | 0.856 | 7.9e-05 | 0.797 | 0.788 | 1.4e-8 |
| WGAN-div + nonlinear *mixup* | 0.853 | 0.858 | 6.5e-05 | 0.792 | 0.784 | 2.2e-8 |
| Standard Sampling | 0.840 | 0.840 | 7.1e-04 | 0.802 | 0.791 | 4.7e-7 |

## Results and discussion

We first compare the classification results of the proposed method, base model (with experimental settings as in Implementation section without oversampling) and other studies conducted on the same dataset, as shown in Table 1. The cell-cycle phases dataset presents a case of extreme data imbalance, where minority classes are in a few tens. In order to handle class imbalance in cell-cycle phases dataset, Eulenberg et al.^1^ used repetition of minority samples, Jin et al.^6^ included WGAN-GP generated images and our previous work^42^ used WGAN-div and *mixup* images with minority focussed sampling approach. The latter approach supplements the mini-batch of original images with another batch of only synthetic samples of minority classes. Furthermore, *mixup* samples were generated from only WGAN-div images. However, in the current study we extend our method^42^ by including nonlinear *mixup* images along with WGAN-div and *mixup* images in the supplementary batch. Additionally, *mixup* and nonlinear *mixup* images are generated from both original and WGAN-div images.

The proposed nonlinear *mixup* approach applies 3D rotation to each pixel of the synthetic image obtained after applying standard mixup on two reference images. In this way, transformations in the spatial domain and colour space are combined to increase the regularisation capability of *mixup*. It is observed that combining transformations in two different domains (spatial and colour) is more advantageous than to further expand the space in the spatial domain to generate synthetic images. We utilised bilinear interpolation, which applies standard *mixup* on the two synthetic images obtained from the standard mixup of the two sets of original images (Fig. 6). The increase in the number of input images expands the space to generate the synthetic image, consequently influencing the regularisation effect. Notably, bilinear interpolation impacted the regularisation effect negatively as the interpolations are applied twice on the original images, which made the final synthetic image drift away from the original pixel distribution. However, this method can help when out-of-distribution sampling is desired.

**Figure 6.**
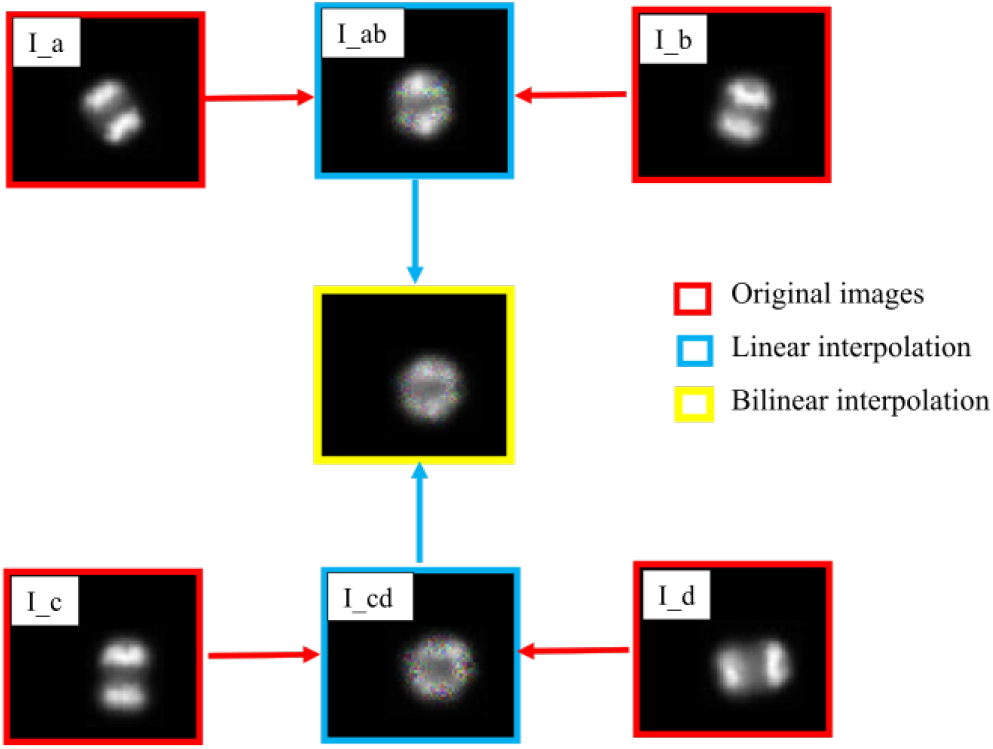
*mixup* using bilinear interpolation.

Data imbalance in the RBC morphologies dataset is medium, however our method demonstrates the performance gain over the existing study^2^ and the base model (Table 1). Doan et al.^2^ applied under-sampling on training and validation datasets, and random combinations of horizontal or vertical flips, horizontal or vertical shifts and rotations to balance the datasets. As the results show, our proposed method achieves superior performance on both datasets compared to the respective state-of-the-art studies. The results also validate the use of more than one approach for data augmentation to create diverse samples for better classification performance. Multiple augmentation approaches help in enhancing the generalisation ability of the trained model and make it perform well on unseen data.

Improvement in individual F1-score of each minority class is demonstrated in Fig. 7. Jin et al.^6^ performed test-time oversampling and Doan et al.^2^ applied test-time under-sampling which could artificially boost the test results, while our results are only from the test images provided in the dataset. Therefore, Fig. 7 compares the per-class F1-scores with other existing studies and respective base models.

**Figure 7.**
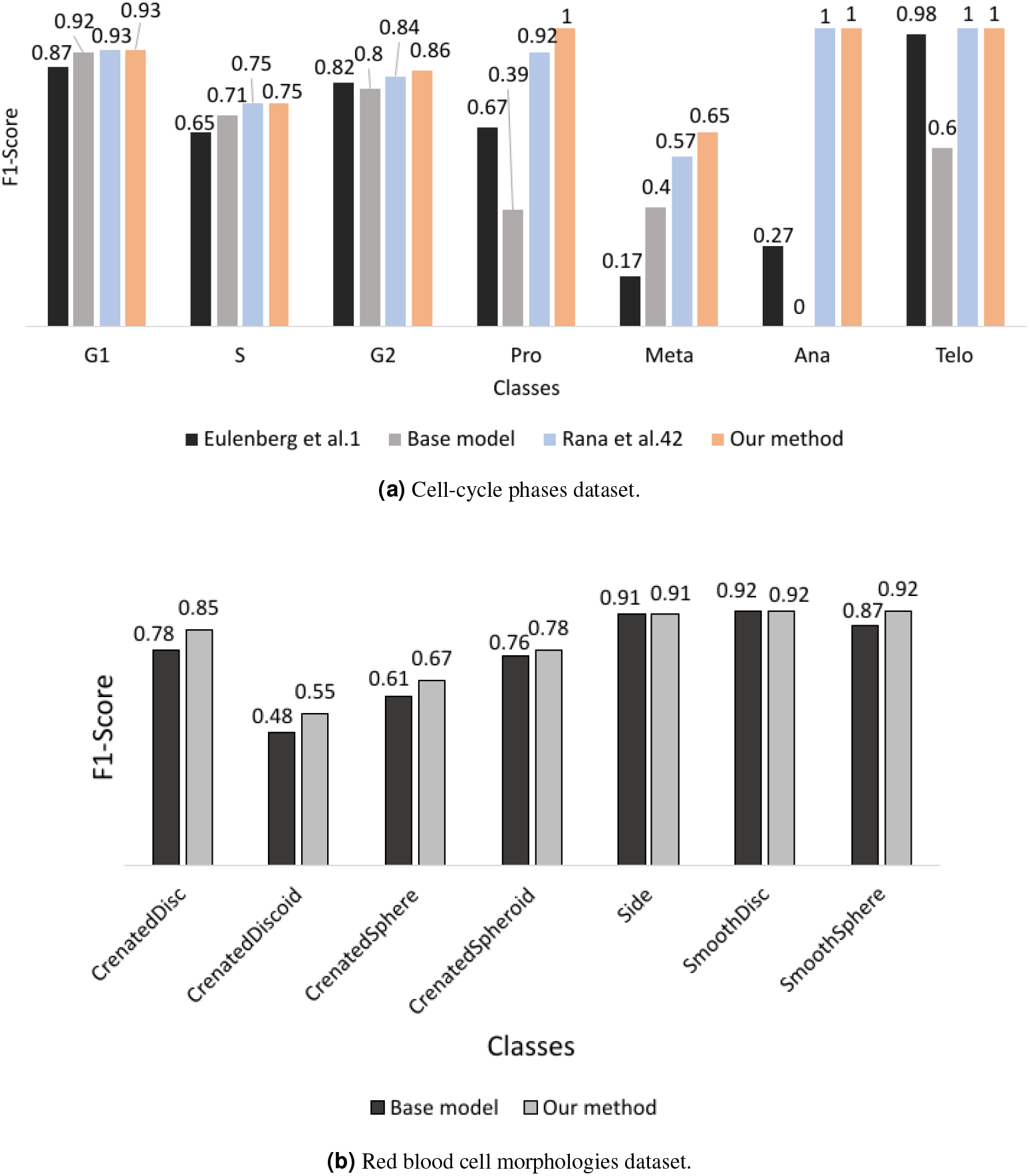
Comparing per-class F1 scores with base model and existing studies on same dataset.

We conducted ablation studies of each component (WGAN-div, *mixup*, nonlinear *mixup* and minority class focussed sampling) of the proposed approach (Table 2). The compared methods are as follows: 1) WGAN-GP + *mixup* + nonlinear *mixup*, which is oversampling with WGAN-GP generated images and corresponding *mixup* and nonlinear *mixup* samples, using minority class focussed sampling; 2) WGAN + *mixup* + nonlinear *mixup*, which is oversampling with WGAN generated images and corresponding *mixup* and nonlinear *mixup* samples, using minority class focussed sampling; 3) GAN + *mixup* + nonlinear *mixup*, which is oversampling with GAN generated images and corresponding *mixup* and nonlinear *mixup* samples, using minority class focussed sampling; 4) WGAN-div + *mixup* + nonlinear *mixup*^12^, *which is oversampling with WGAN-div generated images and corresponding mixup* and nonlinear *mixup* samples using minority class focussed sampling, where nonlinear *mixup* samples were generated following the method presented by Summers et al.^12^; 5) WGAN-div + *mixup*, using only WGAN-div generated images and corresponding *mixup* images for oversampling with the proposed sampling approach; 6) WGAN-div + nonlinear *mixup*, using only WGAN-div generated images and corresponding nonlinear *mixup* images for oversampling with the proposed sampling approach; 7) standard sampling, creating mini-batches using standard random sampling strategy which randomly selects the images from the original and synthetic samples with no supplementary batch as proposed in the current study.

The results from the ablation study (Table 2) demonstrate the performance gain with the proposed framework that includes WGAN-div, *mixup* and nonlinear *mixup*. On comparison with GAN, WGAN and WGAN-GP, the results indicate higher quality of WGAN-div generated samples and exhibit its utility as a data augmentation method for imbalanced classification. To further demonstrate the statistical significance of the results, we performed Wilcoxon rank sum test at significance level of 1%. The Wilcoxon rank sum test compares each ablation study experiment with the proposed method based on the computed probabilities of the correct label of each test set sample. As shown in Table 2, obtained p-values are smaller than 0.01, which implies that the proposed method has statistically more significant performance than the compared methods.

In addition, it is observed that the WGAN-div model demonstrates consistent decrease in the absolute value of critic loss, which converges to comparatively more definite and lower critic loss than other WGAN models (see Supplementary Fig. S1, S2). The curves for WGAN and WGAN-GP show sudden drop to the converged critic loss value, which indicates high probability of overfitting. It is also observed that the Prophase class required higher value of *n*_*critic*_ and lower learning rate than the other classes. It is due to a sub-phase in cell-cycle, namely Prometaphase which is not included as a separate class in the dataset. This sub-phase follows Prophase and precedes Metaphase, and has coincided cellular events with the early Metaphase. Inclusion of Prometaphase images in Prophase has led to higher intra-class difference in the Prophase class and thus requires lower learning rate and more rounds of training to converge as compared to other classes.

Anaphase and Telophase being consecutive phases in cell-cycle have visually similar appearance (Fig. 2(a)). The lower accuracy of Anaphase in previous methods is due to the cells being wrongly classified as Telophase. The obtained results demonstrate the efficacy of oversampling during training with diverse samples for classification when there is low inter-class difference. Likewise, for the RBC morphologies dataset, inaccurate predictions due to similar features in neighbouring classes are reduced with the proposed method.

It is also observed that *mixup* and nonlinear *mixup* achieved similar macro F1-scores for both datasets when used separately, however classification performance is improved if both algorithms are applied together during training. Experiments also show better performance of previously^42^ proposed minority focussed sampling than the standard sampling approach. The proposed sampler finetunes the target distribution and allows the inclusion of the minimum number of synthetic samples required, without affecting the representation of original samples during training. Therefore, it is observed that despite very few samples in Telophase, it still achieved an F1 score of 1 with *n* set to 1. The proposed sampler helps avoid overfitting, reduces the computational cost and does not require a preset sample distribution ratio.

## Conclusion

In this paper, we propose a novel nonlinear *mixup* for data augmentation in image classification problems, and extend our previous work^42^ to demonstrate its superiority in classifying microscopy images of blood cells from two datasets. The *non*-*linear mixup* technique combines transformations in the spatial domain and colour space to further improve the regularisation capability of *mixup*. Nonlinear mixup samples are used for training along with the mixup and WGAN-div generated images through a minority focussed sampling approach. Our experimental results demonstrate the advantages of using three different data augmentation approaches for oversampling of minority classes to increase the diversity of the training dataset for more robust classification. The proposed classification pipeline outperforms state-of-the art approaches on publicly available cell-cycle phases dataset and red blood cells morphologies dataset.

## Data availability

Dataset for annotated images of different phases of cell-cycle is publicly available at https://bbbc.broadinstitute.org/BBBC048. Dataset for annotated images of different phenotypes of red blood cells is publicly available at https://figshare.com/articles/URL7_Annotated_Data/12432506.

## Acknowledgements

This research was undertaken with the assistance of resources and services from the National Computational Infrastructure (NCI), which is supported by the Australian Government. This research is supported by an Australian Government Research Training Program Scholarship.

## Author contributions statement

All authors contributed to conceptualisation of the study. P.R. and Y.S. designed the methodology. P.R. performed the experiments, analysed the results and wrote the manuscript. Y.S., E.M. and A.S. reviewed the draft, provided inputs at various stages and contributed to the writing of the manuscript. All authors read and approved the final manuscript.

