## Supplementary materials for "Data augmentation for imbalanced blood cell image classification"

### Supplementary Information

**Supplementary Figure S1:** Critic loss - epoch training curves for cell-cycle phases dataset.

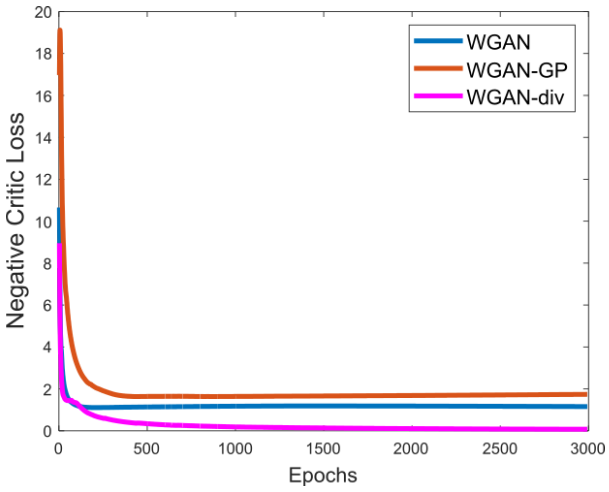


**(a)** Anaphase.

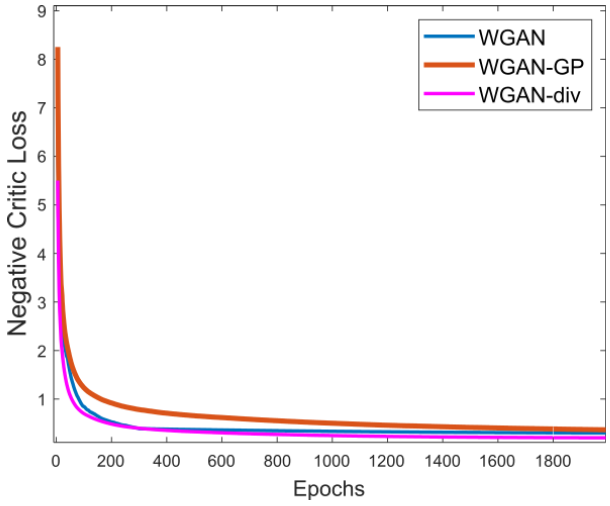


**(b)** Metaphase.

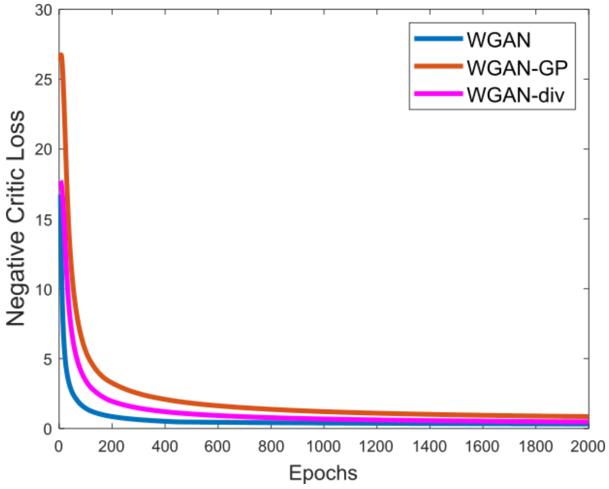


**(c)** Prophase.

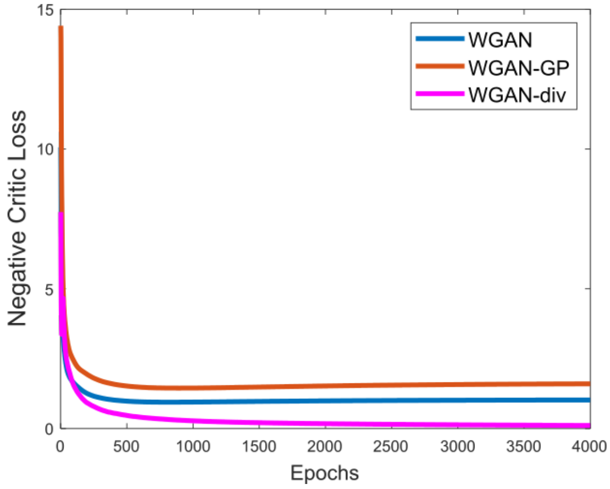


**(d)** Telophase.

**Supplementary Figure S2:** Critic loss - epoch training curves for RBC morphologies dataset.

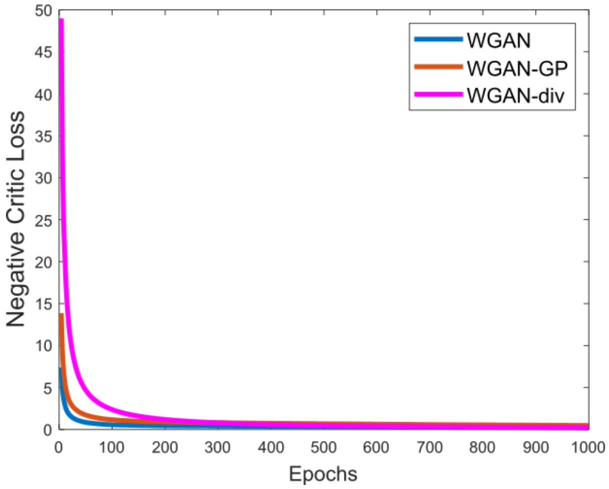


**(a)** CrenatedDiscoid.

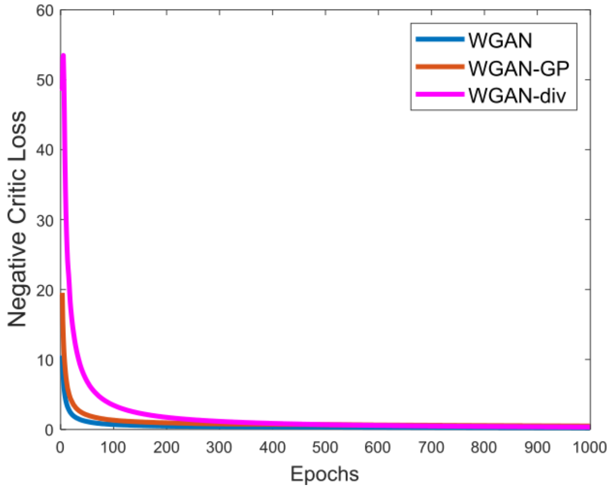


**(b)** CrenatedSphere.

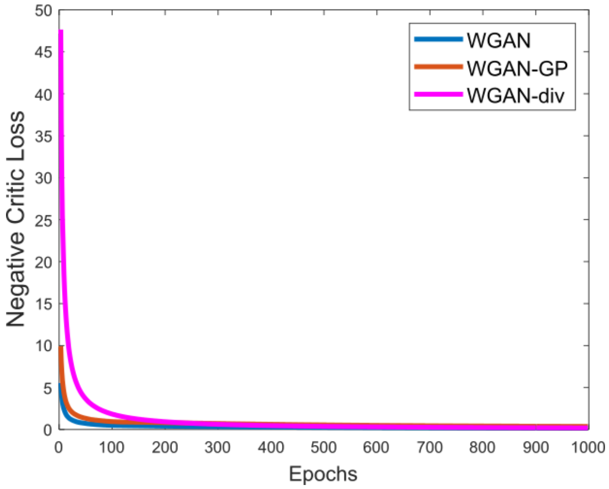


**(c)** CrenatedSpheroid.

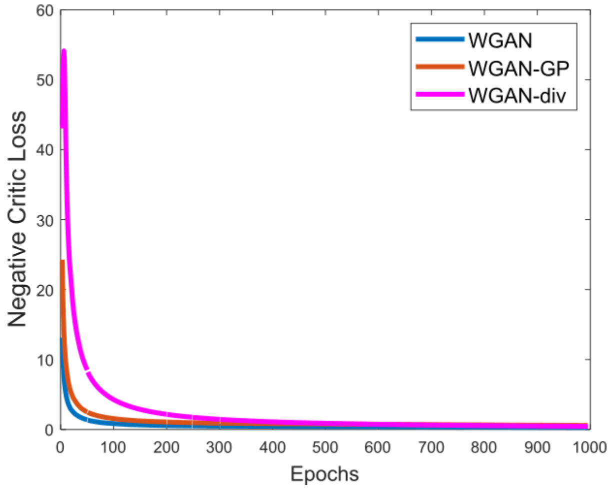


**(d)** SmoothSphere.

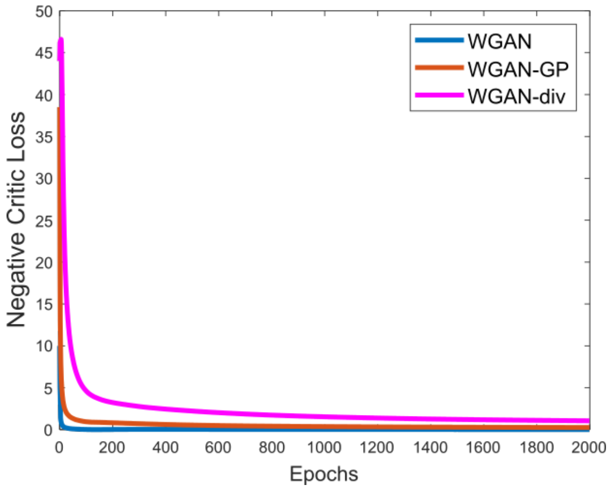


**(e)** CrenatedDisc.

**Supplementary Figure S3:** FID Score vs epoch for Cell-cycle phases dataset.

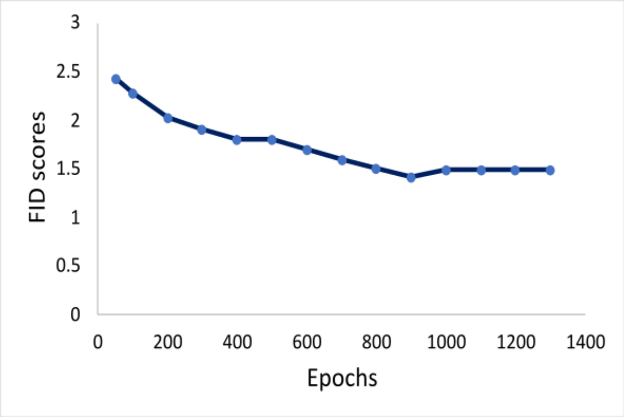


**(a)** Anaphase.

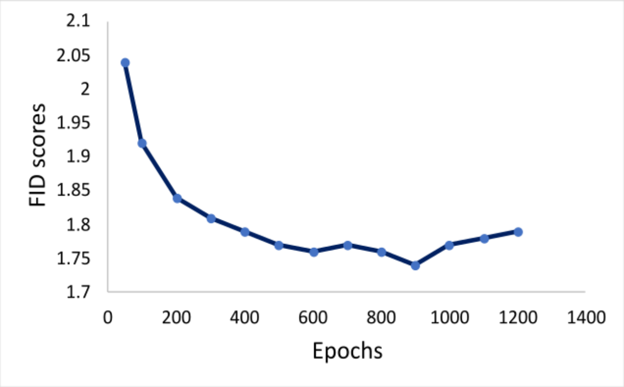


**(b)** Prophase.

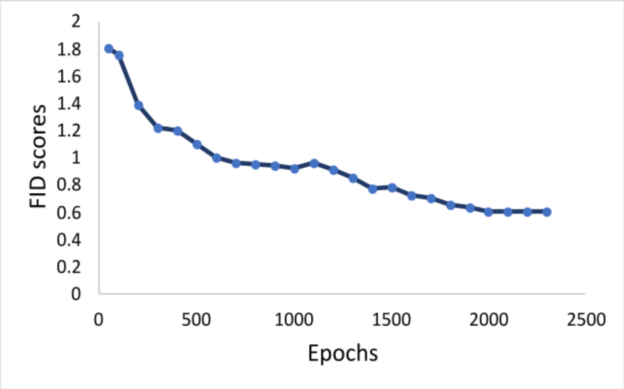


**(c)** Metaphase.

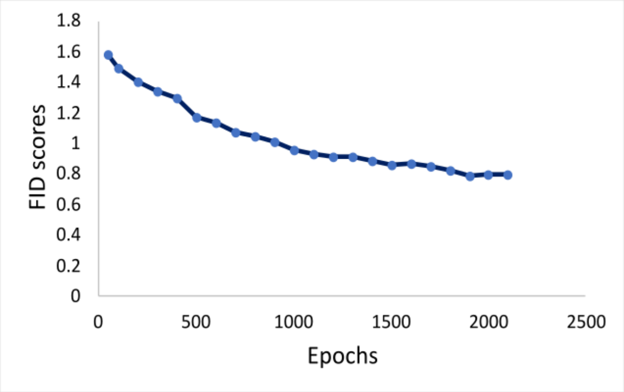


**(d)** Telophase.

**Supplementary Figure S4:** FID Score vs epochs for RBC morphologies dataset.

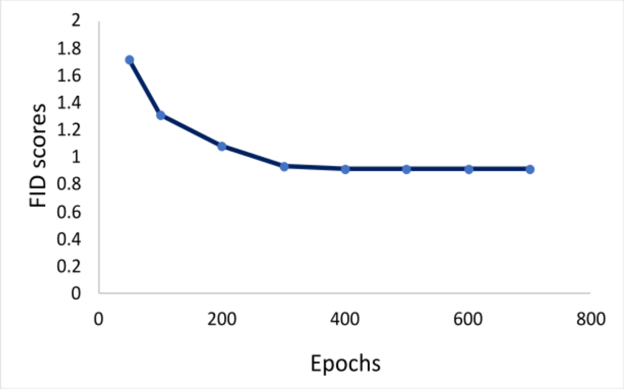


**(a)** CrenatedDiscoid.

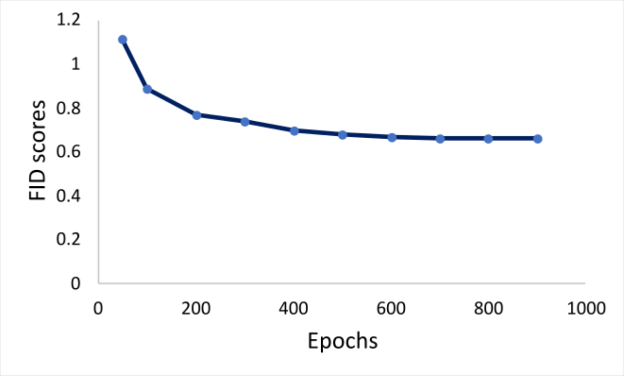


**(b)** CrenatedSphere.

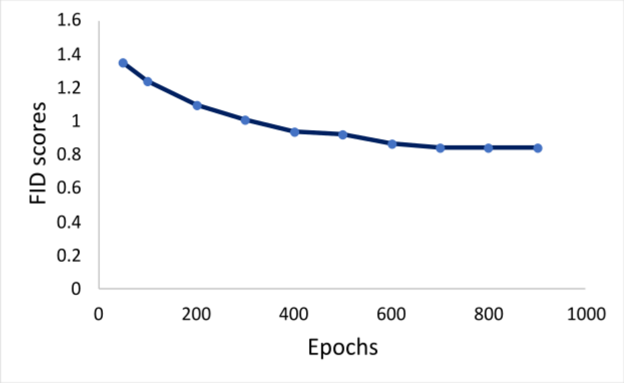


**(c)** CrenatedSpheroid.

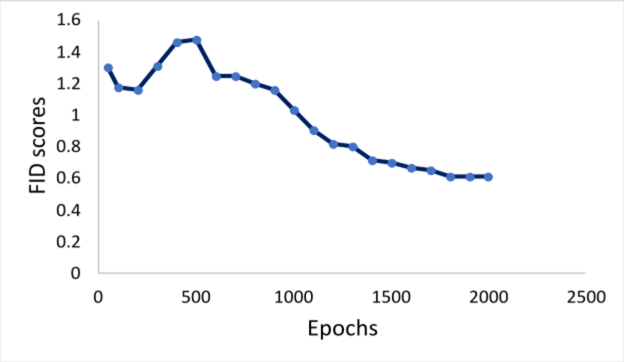


**(d)** SmoothSphere.

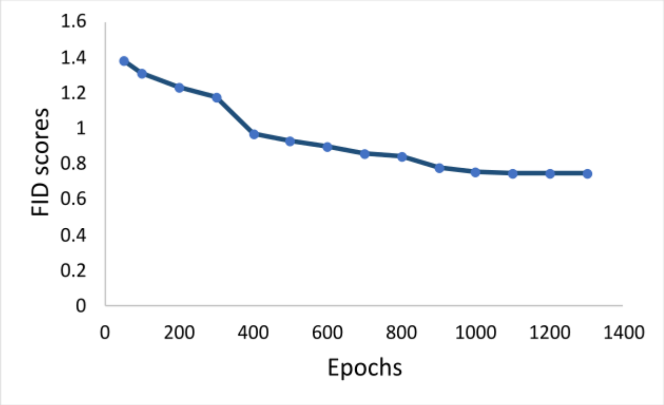


**(e)** CrenatedDisc.
